# Activation of G-protein coupled estradiol receptor 1 in the dorsolateral striatum attenuates preference for cocaine and saccharin in male but not female rats

**DOI:** 10.1101/824078

**Authors:** Jacqueline A. Quigley, Jill B. Becker

## Abstract

There are sex differences in the response to psychomotor stimulants, where females exhibit a greater response than males, due to the presence of the gonadal hormone estradiol (E2). Extensive research has shown that E2 enhances drug-seeking and the rewarding properties of cocaine for females. The role of E2 in male drug-seeking, however, is not well understood. The current study investigated pharmacological manipulation of E2 receptors in the dorsolateral striatum (DLS) on preference for cocaine in gonad-intact male and female rats. In males, activation of G-protein coupled E2 receptor 1 (GPER1), via administration of ICI 182,780 or G1, attenuated conditioned place preference for 10mg/kg cocaine, while inhibition of GPER1, via G15, enhanced preference at a 5mg/kg cocaine dose. Similarly, GPER1 activation, via G1, prevented males from forming a preference for 0.1% saccharin (SACC) versus plain water. Surprisingly, activation of GPER1 did not alter preference for cocaine or SACC in females. These studies also examined the quantity of E2 receptor mRNA in the dorsal striatum, using qPCR. No sex differences in relative mRNA expression of ERα, ERβ, and GPER1 were observed. However, there was greater GPER1 mRNA, relative to ERα and ERβ, in both males and females. The results presented here indicate that E2, acting via GPER1, may be protective against drug preference in male rats.

## Introduction

More men report abusing cocaine and qualify for cocaine use disorder, however, cocaine use in women is gradually increasing and there is growing evidence to suggest that women are more vulnerable to addiction ^1^. The time of first drug use to admission for addiction treatment is typically shorter for women than men ^2^. When entering treatment, women also present with greater social, behavioral, and psychological symptoms related to substance use disorder, despite having abused drugs for a shorter period of time ^3,4^. These women are also taking greater amount of drug when entering treatment and report experiencing enhanced cravings compared to their male counterparts ^5,6^. Historically, we have attributed sex-specific behaviors related to addiction as being mediated by cultural influences combined with differences in neurobiological function between males and females, including a sex-specific role of the gonadal hormone estradiol (E2) that enhances drug-taking in females ^7,8^.

Research into the biological bases for sex differences in cocaine addiction has focused on how E2 alters the rewarding properties of cocaine in females. Clinical models have found that when E2 levels are high, women report an enhanced euphoria or “high” when abusing smoked cocaine ^9^. Similarly, female rats show enhanced cocaine craving and motivation for cocaine when E2 levels are highest ^10–13^. While these data suggest that E2 plays a major role in facilitating motivation and other behaviors in females, the role of E2 in modulating males’ addiction-like behaviors should also be understood. Testosterone produced by the male testes is aromatized to E2 in the brain and periphery to act on E2 receptors in the male brain.

E2 acts by binding to receptors alpha (ERα), beta (ERβ), and G-protein coupled E2 receptor 1 (GPER1). The cellular location of traditional E2 receptors α and β are either in the cell nucleus, or after palmitoylation, on the extracellular membrane in association with caveolin proteins ^14–16^. GPER1 activates intracellular signaling cascades mediated by cAMP, ERK, and PI3K and localized on the plasma membrane and in the endoplasmic reticulum ^17^. In females, ERα and GPER1 in the dorsal striatum are localized to GABAergic medium spiny neurons with recurrent collaterals onto dopamine (DA) terminals, as well as cholinergic interneurons and glia ^18,19^. The localization of ERα and GPER1 in the dorsal striatum of males remains unknown.

For males and females alike, the rewarding effects of cocaine are attributed to the drug’s direct effects on DA reuptake in the striatum ^20,21^. However, there are sex differences in the effects of cocaine on increases in extracellular DA and receptor binding that are driven by E2 in the dorsal striatum. In females, E2 binding to GABAergic neurons decreases GABA release and this disinhibits DA terminals and ultimately increases stimulated DA release locally ^16,22^. This E2-induced increase in DA release is associated with an enhancement of the effects of cocaine and other psychostimulants on drug-induced behaviors, such as behavioral sensitization ^23–26^. Previous research has not found that E2 has similar effects on cocaine-induced increases in DA in males ^23^. Finally, the effects of E2 in dorsolateral striatum (DLS) are mediated by mGluR signaling in females ^24^

There is extensive support for E2 in mediating drug abuse liability in females but a lack of attention to understanding the role of E2 in males. There is recent evidence from gene knockout studies, however, that GPER1 could be playing a modulatory role in the preference for morphine in males ^27^. Given that previous research has highly implicated the DLS as an important region for studying the effects of E2 on addiction-like behaviors in females, the current set of studies were deigned to investigate how intra-DLS GPER1 modulates preference for rewarding stimuli in males as well as females. A cocaine conditioned place preference (CPP) and saccharin (SACC) two-bottle choice behavioral paradigms were used to assess how GPER1 activation or inhibition alters preference for rewarding stimuli. Finally, the current study also used qPCR to determine relative mRNA levels of ERs in the dorsal striatum of both males and females.

## Materials and Methods

### Animals

A total of 62 male and 46 female gonad-intact Sprague-Dawley rats were used in the current set of experiments, as detailed in Figure 1A. Animals were ordered from Charles River Breeding Laboratory (Portage, MI, USA) and were approximately 75 days old on arrival. Animals were maintained on a 14:10 light/dark cycle in a temperature-controlled climate of 72°F ± 2°F, in ventilated laboratory cages. Rats had ad libitum access to water and phytoestrogen-free rat chow (2017 Teklad Global, 14% protein rodent maintenance diet, Harlan rat chow; Harlan Teklad, Madison, WI, USA). Animals were initially housed in same-sex pairs until undergoing surgery, after which they were subsequently housed individually. All animals were weighed daily to determine good health and at this time, females were also vaginally lavaged to determine stage of estrus. All animal care and experimental procedures were carried out in accordance with the National Institutes of Health guidelines on laboratory animal use and care, using a protocol approved by University of Michigan Institutional Use and Care of Animals Committee.

**Figure 1.**
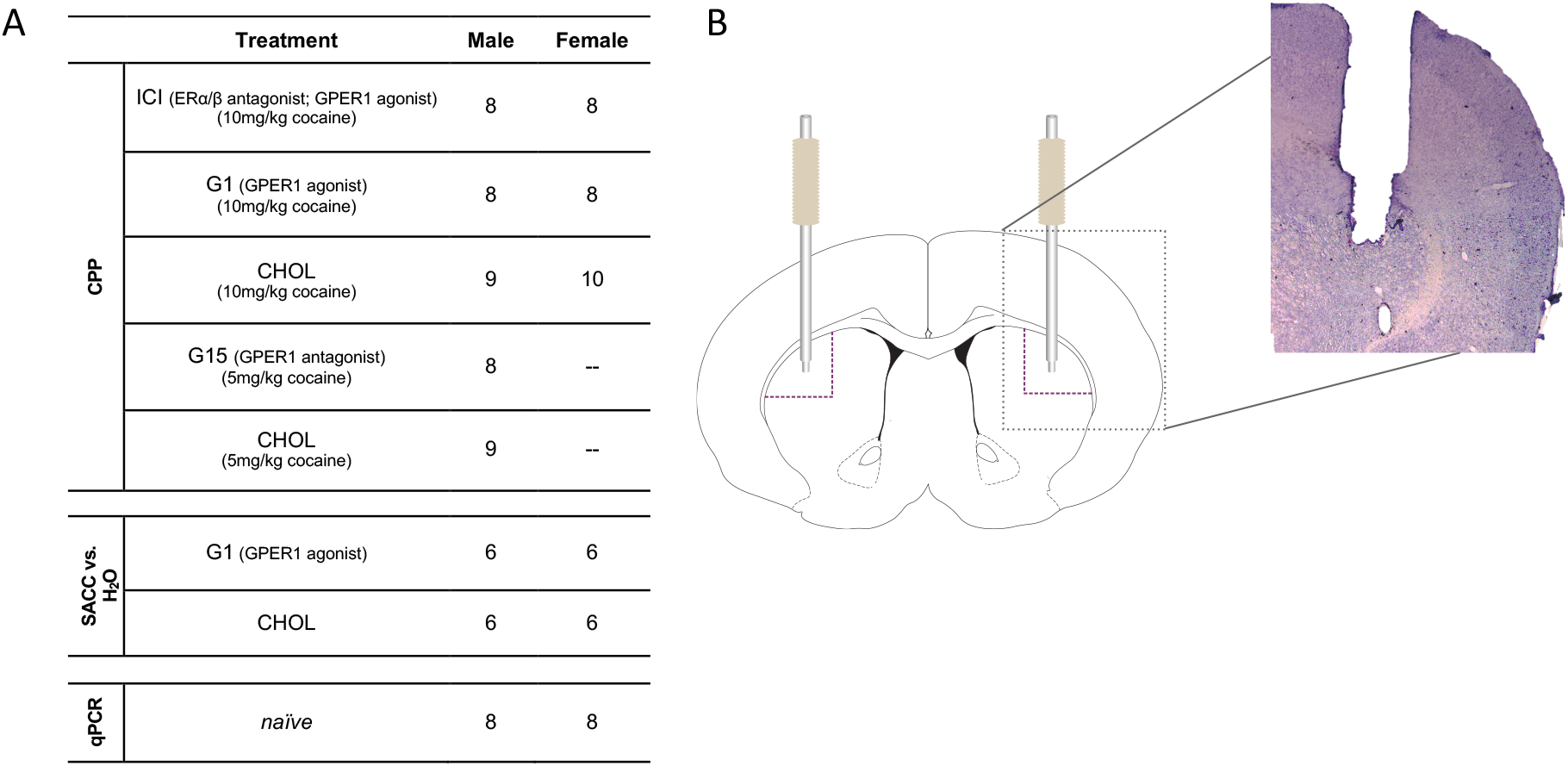
(A) Number of male and female animals per treatment condition for each experiment. **(B)** Representative cannula placement into the DLS.

### Stereotaxic Surgery and Drug Preparation

One week after arriving in the laboratory, rats underwent surgery for the implantation of bi-lateral guide cannula aimed at the DLS (AP: +0.4 ML: +/-3.6 DV: -4.0). On the day of surgery, rats were injected with carprofen (5mg/kg s.c.) and 30 minutes later were anesthetized with ketamine (50mg/kg i.p.) and dexmedetomidine (0.25mg/kg i.p.), then prepared in a stereotaxic frame. At the conclusion of the surgery, rats were given atipamezole hydrochloride (0.5mg/kg i.p.) and 3ml 0.9% saline (s.c.). Every 24 hours for three days post-surgery, rats were given carprofen (5mg/kg s.c.) prophylactically for post-operative pain. No animal underwent behavioral testing for at least 7 days after surgery.

During surgery, 33-gauge solid stylets were inserted into the 26-gauge hollow guide cannula that were fixed on animals’ skull. These stylets were flush with the bottom of the guide cannula and did not protrude into the brain. Treatment conditions were randomly assigned to animals prior to behavioral testing. Control animals received 100% cholesterol (CHOL) and experimental animals received either 10% ICI (ERα/ERβ antagonist; GPER1 agonist), G1 (agonist targeting GPER1) or G15 (antagonist targeting GPER1) dissolved in Cholesterol (Control) via stylets, which protruded from the guide cannula by 1mm and delivered treatment directly into the DLS. Treatment stylets were prepared as previously described ^28^. In order to insert stylets, rats were briefly anesthetized with 5% isoflurane.

Hollow guides and interlocking treatment stylets were manufactured by and purchase from P1 Technologies. Drugs were obtained from the following sources: ICI182,780 (ICI) (Santa Cruz Biotechnology, purity ≥ 98%); G1 (Cayman Chemical, purity ≥ 98%); G15 (Cayman Chemical, purity ≥ 95%); Cholesterol (Santa Cruz Biotechnology, purity ≥ 92%)

### Conditioned Place Preference (CPP)

Animals were tested on a CPP paradigm that took place over 10 consecutive days, as illustrated in Figure 2 A. The CPP apparatus consisted of two side chambers (15.5 inches x 12 inches) and a center neutral chamber (15.5 inches x 7.5 inches). On day 1 (pre-test), rats were placed in the novel chamber and were allowed to move freely between all compartments for 30 minutes. Immediately following pre-test session, treatment stylets were inserted. For eight days thereafter, animals were trained to associate each of the three chambers with another stimulus (drug paired; neutral; vehicle paired). Each conditioning session began with a 10-minute habituation in the center neutral chamber. Animals were then removed from the chamber and received an intra-peritoneal (i.p.) injection of either cocaine or vehicle and immediately placed in one of the side chambers (drug paired; vehicle paired) for 30 minutes. Animals were conditioned to each stimulus (drug or vehicle) 4 times each, every other day. Which treatment animals received first was counterbalanced. On day 10 (Test), rats were placed in the three-compartment chamber and were again allowed to move freely between all compartments for 30 minutes.

**Figure 2.**
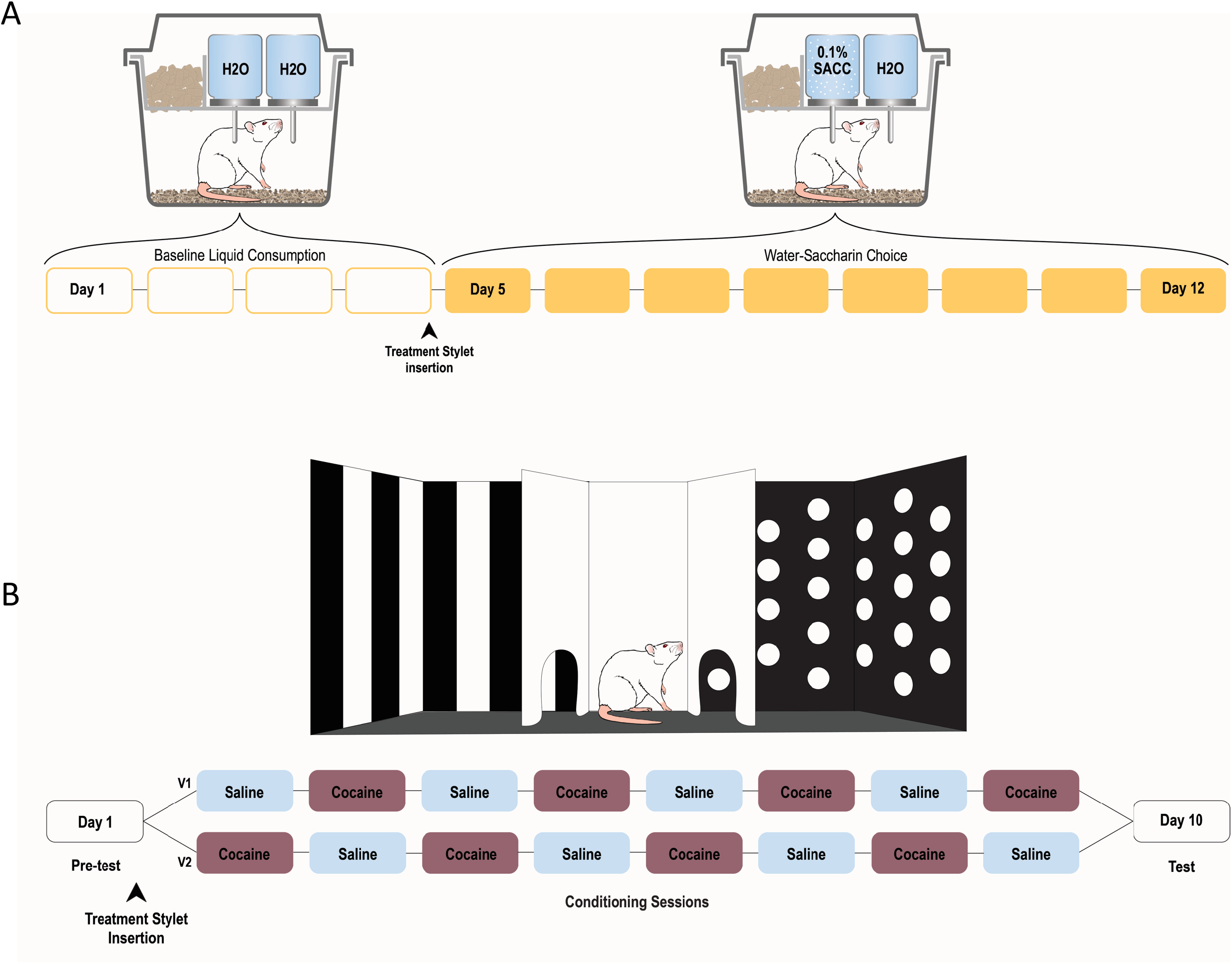
(A) The apparatus used for conditioned place preference training and testing was a three-compartment chamber with three areas, differentiated by distinct tactile and visual cues. As described in the timeline, the pre-test took place on day 1, followed by 8 conditioning sessions, ending with a final test session. Version 1 (V1) and version 2 (V2) refer to conditioning either beginning with saline or cocaine on day 2, which was counterbalanced across animals. Treatment stylets were inserted after pre-test on day 1 and remained inserted for continuous treatment through the final test session. **(B)** A two-bottle choice paradigm was utilized to determine preference for 0.1% SACC versus plain water. Two bottles were accessible to individually housed animals in their home cages. On days 1-4, both bottles contained H_2_O only. During days 5-12, one bottle contained H_2_O and the other contained 0.1% SACC dissolved in H_2_O. Treatment stylets were inserted before SACC introduction on day 5 and remained for the duration of the experiment.

The side in which animals spent the most time during pre-test was treated as their “preferred chamber”, which differed for each animal. A biased design was utilized, so that each individual animal’s initially preferred chamber was paired with saline, and their initially un-preferred chamber was paired with cocaine for conditioning. ANYMAZE tracking software (Stoelting Co., Wood Dale, IL 60191) was utilized to track the amount of time spent in each chamber.

Either a 5mg/kg or 10mg/kg cocaine dose was utilized during conditioning in order to be able to test both an increase and decrease in cocaine CPP after ER manipulation. For example, both males and females acquire cocaine CPP at a 10mg/kg conditioning dose therefore we utilized that dose when investigating a potential decrease in CPP (i.e., ICI and G1). When investigating whether there was an increase cocaine CPP in males, we used a dose that did not generally produce a CPP, 5mg/kg, which allowed us to identify a potential increase in CPP (i.e., G15) without a ceiling effect. This dose of cocaine did produce CPP for cocaine in females (data not shown), so we did not test the effects of G15 in females with 5mg/kg cocaine.

### Two Bottle Choice Experiment (SACC versus H_2_O)

Animals were tested on a two-bottle choice paradigm that took place over 12 consecutive days, as illustrated in Figure 1 B. Briefly, on days 1-4, two bottles both containing water were available on each animal’s home cage to determine that were was no significant difference between total liquid intake between animals. During days 5-12, one bottle contained water and the second bottle contained 0.1% saccharin (Sigma-Aldrich; purity ≥ 92%) dissolved in water. Placement of the bottle (left versus right) was switched daily to account for a potential side of cage preference. On day 5, four hours before the SACC bottle was introduced, stylets containing either 10% G1 in cholesterol or cholesterol alone were inserted and remained in place for the duration of the experiment. Once daily, one hour prior to the start of dark cycle, bottles were removed from home cages, weighed, and refilled.

### Euthanasia and Tissue Preparation

Animals received 0.5 ml of Sodium Pentobarbital (i.p). Once the animal was fully sedated, it was perfused transcardially with 0.1M phosphate buffered saline followed by 4% paraformaldehyde. The brain of each rat was also dissected and post-fixed in 4% paraformaldehyde for 24 hours and afterwards stored in 10% sucrose. Brains were sliced on either a microtome or cryostat in 60-micron sections then were mounted on slides, stained with cresyl violate and cover slipped. Sections were analyzed for accurate guide cannulae placements, depicted in Table 1, by an observer blind to experimental conditions. Only animals that had accurate cannulae placements are shown and were included in the final analysis. For animals in the H_2_O versus SACC experiment, females’ ovaries and uterus and males’ testes and vas deferens were dissected and weighed after the animal was perfused as a proxy to determine if G1 treatment in the brain affected peripheral gonadal tissues.

### Quantitative Polymerase Chain Reaction (qPCR)

Naïve gonad-intact male and female rats were given 0.5 ml of FatalPlus (i.p.; 195 mg sodium pentobarbital; Vortech Pharmaceuticals, Ltd; Dearborn, MI). Once the animal was fully anesthetized the brain was rapidly removed and placed into ice-cold saline. The dorsal striatum from both hemispheres was micro-dissected from each animal and stored at -80°C for later processing.

Tissue was extracted using a phenol-chloroform reaction using Trizol (Cat. No. 97064-950, Amresco) as the lysis reagent. Next, a QuantiTect® Reverse Transcription Kit was used for genomic DNA wipeout and cDNA synthesis (Cat. No. 205314, Qiagen). Relative gene expression was measured using a RealMasterMix™ Fast SYBR Kit (Cat. No. A25742, Applied Biosystems). Primers (not designed to determine specific splice variants, but to detect all variants) were purchased from Qiagen for: ERα (Cat. No. QT00386925, Qiagen Primer; reference sequence NM_012689), ERβ (Cat. No. QT00190113, Qiagen Primer; reference sequence NM_012754), and GPER1 (Cat. No. QT00376943, Qiagen Primer; reference sequence NM_133573). These genes were compared against the housekeeping gene HPRT1 (Cat No. QT00199640, Qiagen Primer) and relative gene expression was quantified using the 2^ddCT method. Samples were run in triplicates at the Biomedical Research Core Facilities at the University of Michigan (https://brcf.medicine.umich.edu/cores/advanced-genomics/technologies/real-time-pcr/).

### Statistical Analysis

CPP data were analyzed using 2-way ANOVAs. In the case of a significant interaction, a Bonferroni correction was used for multiple comparisons. For males and females independently, CPP data were analyzed by time spent in the drug-paired chamber (pre-test versus test) between treatment conditions, within each sex. Behavioral testing for males and females was not done simultaneously therefore, we did not compare them statistically.

Daily preference of 0.1% SACC versus water was calculated as a percentage: (0.1% SACC consumed/(0.1%SACC + Water consumed))*100. An unpaired t-test was performed to identify treatment group differences in preference score. Unpaired t-tests were performed to determine if gonad weights were different between G1 or CHOL treated males or females.

Relative mRNA expression of ERα, ERβ, and GPER1 was analyzed between sexes. Unpaired t tests were performed to identify sex differences in ERβ and GPER1 expression within the dorsal striatum. Nonparametric testing was conducted for ERα because data were not normally distributed for males.

All statistical analyses were performed using GraphPad Prism v8.0 and IBM SPSS Statistics v27.0. All data sets were tested for normality and equal variance between groups. Effect sizes are reported as partial eta (n^2)^ or partial eta-squared (n^2^p) for F-tests and Cohen’s d (d) for t-tests. The threshold for significance in all experiments was set to p<0.05 and sample sizes per experiment were determined based on pilot studies. Animals were excluded from statistical analyses if the placements of the guide cannula were off target in both the CPP and SACC versus H_2_O experiments (< 5% of total animals were excluded).

## Results

### Effects of ER manipulation on cocaine CPP in males

The effects of ER manipulation on cocaine CPP in males, as measured by time spent in the drug-paired chamber, is shown in Figure 3 A-C. Administration of ICI, a nonselective ERα/β antagonist and GPER1 agonist, attenuated males’ preference for 10mg/kg cocaine (Figure 3A). A two-way repeated measures ANOVA revealed a treatment x test session interaction (F _(1,15)_ = 5.758; p = 0.0299, n^2^p = 0.277). A Bonferroni multiple comparisons test revealed that CHOL treated animals increased time spent in the cocaine-paired chamber (4.745 ± 0.974, p = 0.0004) but ICI treated males did not (0.482 ± 1.033, p > 0.9999).

**Figure 3.**
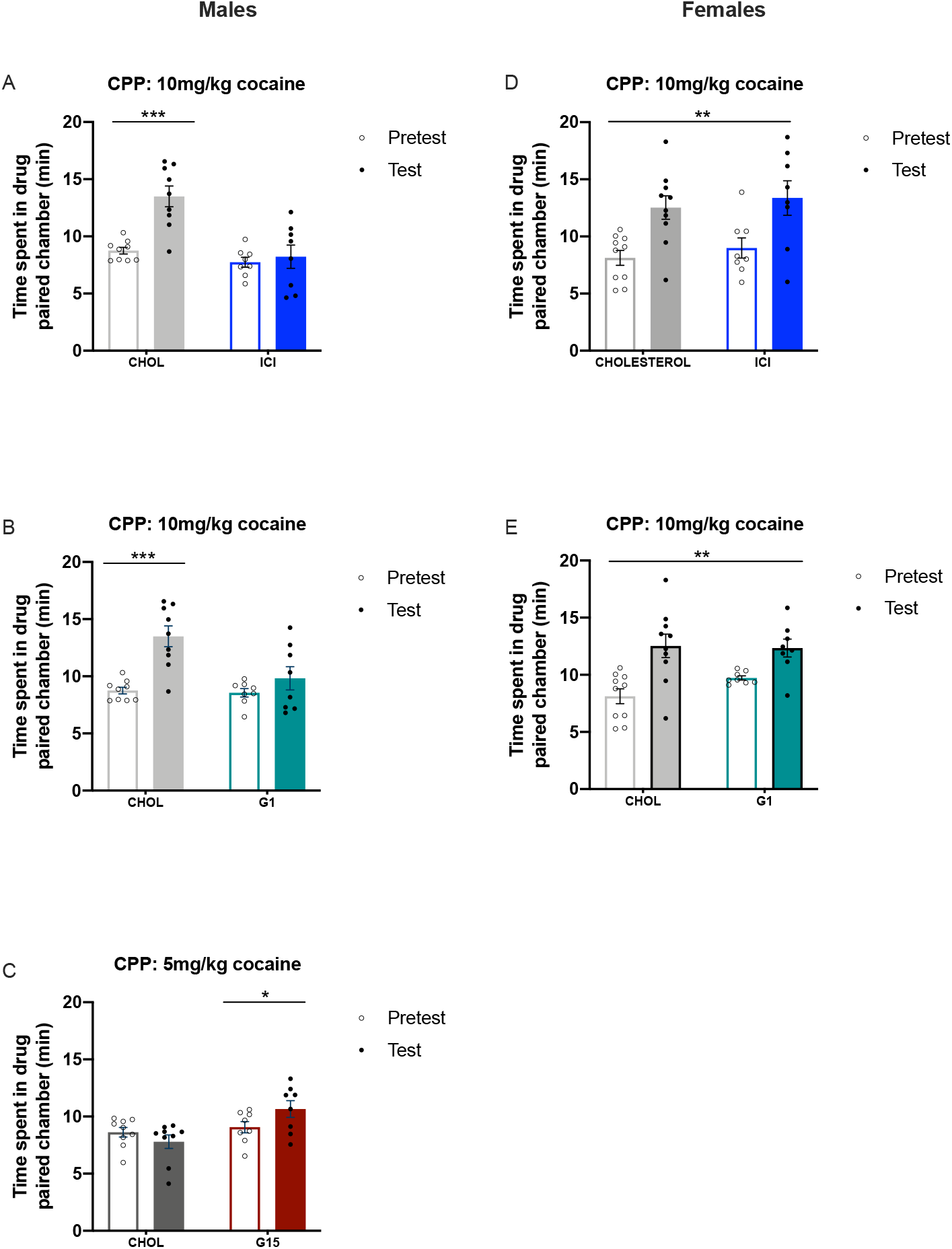
GPER1 activation/inhibition intra-DLS alters cocaine CPP for males but not females. A cocaine CPP is inferred if animals spend more time in drug-paired chamber during test versus pre-test. **(A)** At a conditioning dose of 10mg/kg cocaine, CHOL (control) treated males form a CPP however, treatment of ICI (nonselective ERα/ERβ antagonist; GPER1 agonist) or **(B)** G1 (selective GPER1 agonist) attenuates males CPP for cocaine. **(C)** Treatment of G15 (selective GPER1 antagonist) causes a CPP for a 5mg/kg dose of cocaine; CHOL treated males do not form a CPP. **(D-E)** Females treated with CHOL, ICI, and G1 form a CPP for 10mg/kg cocaine. CPP data are shown as mean +/-SEM and each point corresponds to an individual animal. *p < 0.05, **p < 0.01, ***p < 0.001.

The goal of our follow up study was to differentiate between the effects of ICI on inhibition of ERα and ERβ versus activation of GPER1. We did this by using G1, a selective GPER1 agonist. Figure 3 B represents males treated with G1 compared to CHOL. A two-way repeated measures ANOVA revealed that G1 treatment replicated the behavioral effect of ICI treatment on cocaine CPP; there was a significant interaction between test session and treatment (F _(1,15)_ = 7.429; p = 0.016, n^2^p = 0.331) and multiple comparisons revealed that at 10mg/kg cocaine, G1 treated males did not acquire a CPP for cocaine (1.272 ± 0.972, p = 0.3804), but CHOL animals did (4.745 ± 0.8741, p = 0.0001).

We further investigated whether administration of the GPER1 antagonist, G15, could cause enhanced preference for cocaine in males. Illustrated in Figure 3 C, at a 5mg/kg conditioning dose, CHOL treated males did not show a CPP for cocaine however, G15 treated males did. A two-way repeated measures ANOVA revealed a significant interaction between test session and treatment (F _(1,15)_ = 8.194; p = 0.0119, n^2^p = 0.353). A Bonferroni multiple comparisons test revealed that males treated with G15 spent more time in the drug-paired chamber after conditioning while CHOL treated males did not (1.591 ± 0.6143, p = 0.0406; 0.826 ± 0.579, p = 0.3182; respectively).

### Effects of ER manipulation on cocaine CPP in females

The effects of ER manipulation on cocaine CPP in females are shown in Figure 3 D-E. When cocaine CPP was determined in females treated with CHOL, ICI, or G1, all groups formed a preference for 10mg/kg cocaine. A two-way repeated measures ANOVA was performed to compare test sessions by treatment. Figure 3 D represents CHOL and ICI treated females, where there was a main effect of test session (F _(1,16)_ = 13.99; p = 0.018, n^2^p = 0.467). There was no main effect of treatment however, and no interaction between test session and treatment. Similar results are shown in Figure 3 E, which represents CHOL and G1 treated females, where there was only a main effect of test session (F _(1,16)_ = 20.05; p = 0.0004, n^2^p = 0.556).

### Effects of intra DLS ER manipulation on saccharin preference in males and females

The effects of the GPER1 agonist, G1, on preference for 0.1% SACC versus water is illustrated in Figure 3 D. An unpaired t-test comparing preference scores between CHOL and G1 treated males revealed significant group differences (t _(10)_ = 2.589; p = 0.0270, d = 1.495) in preference scores. There were no differences in total water intake on days 1-4, prior to treatment. There were also no significant differences in total liquid consumed on days 5-12 between treatment groups.

Consistent with the effects of G1 on cocaine CPP, there was no effect of G1 on females’ preference for 0.1% SACC over water, shown in Figure 4 C. An unpaired t-test was used to reveal no significant group differences (t _(10)_ = 0.7998; p = 0.4424, d = 0.462). Both G1 and CHOL groups had equal variance and data were normally distributed. There were no differences in total liquid consumption on days 1-4, prior to treatment or during treatment on days 5-12.

**Figure 4.**
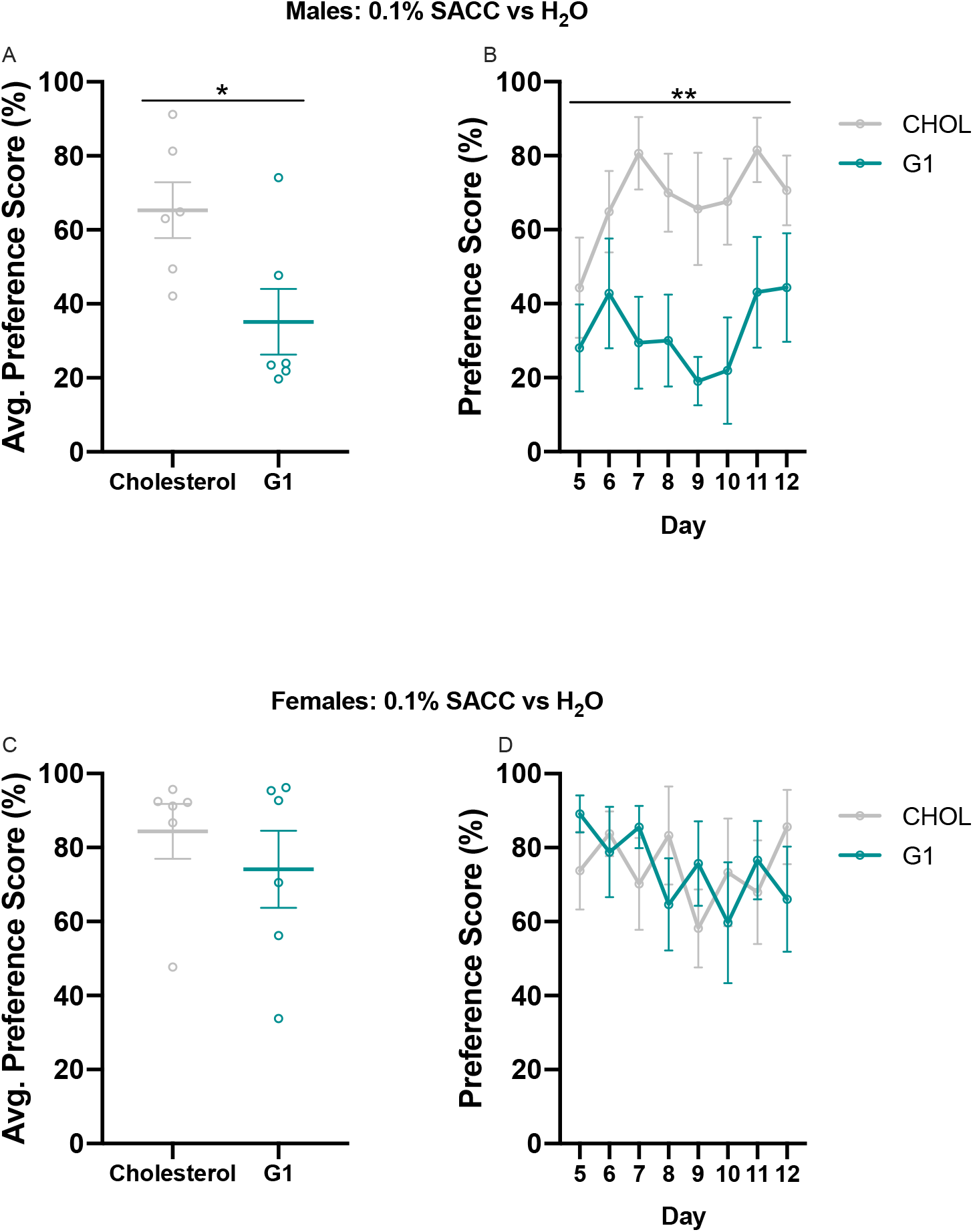
GPER1 activation intra-DLS attenuates preference for 0.1% SACC versus H_2_O for males but not females. **(A)** G1 treatment causes a conditioned avoidance of 0.1% SACC, as indicated by a preference score < 50%. G1 treated males have a significantly lower preference score, averaged across days, than CHOL (control) males. **(B)** Both G1 and CHOL (control) treated females similarly formed a preference for 0.1% SACC over H_2_O, as indicated by group preference scores exceeding 50%. Preference score data are shown as mean +/-SEM with data points representing individual days. *p < 0.05, **p < 0.01.

### Effects of intra DLS ER manipulation on gonad weights in males and females

There was no effect of G1 treatment intra-DLS on gonad weight in either males or females. In females, this included uterus weight (t _(10)_ = 0.4571; p = 0.6574, d = 0.263) and ovary weights (t _(10)_ = 1.892; p = 0.0878. d = 1.09). For males, this included testes (t _(10)_ = 0.1270; p = 0.9014, d = 0.073) and vas deferens weights (t _(10)_ = 1.625; p = 0.1352, d = 0.938).

### Relative mRNA expression of ERα, Erβ, and GPER1 in the dorsal striatum of males and females

As shown in Figure 5 A-C, there were no sex differences in relative mRNA expression of ERs in the dorsal striatum. For ERα, male data points violated normality testing, therefore an unpaired nonparametric Mann-Whitney U test was performed to identify group differences. No sex differences in ERα expression were found (U = 19; p = 0.1949, n = 0.142). Unpaired t tests revealed no group differences for ERβ (t _(14)_ = 0.3334; p = 0.7438, d = 0.166), or GPER1(t _(14)_ = 0.7504; p = 0.4654, d = 0.374).

**Figure 5.**
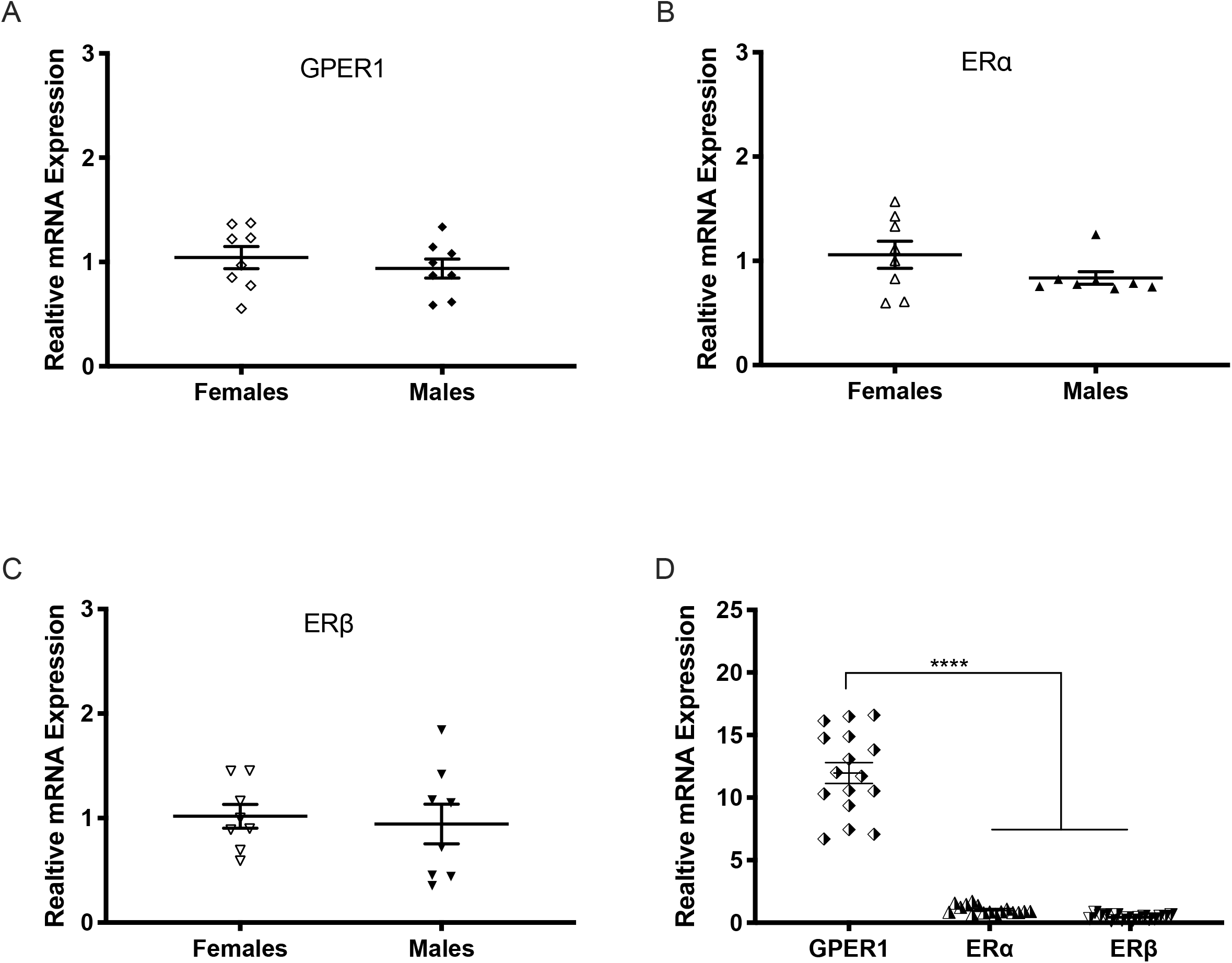
(A-C) There are no significant differences in relative mRNA expression of estradiol receptors in the dorsal striatum between females and males. Data are shown as males relative to females. **(D)** Relative mRNA expression of GPER1 is greater than ERα and ERβ, which do not differ from one another. Data are shown as GPER1 and ERβ relative to ERα. Data are shown as mean +/-SEM with data points representing individual animals. ****p < 0.0001

The expression of GPER1 mRNA was greater than ERα and Erβ mRNA relative to the HRPT housekeeping gene, as illustrated in Figure 5D. A significant one-way ANOVA determined differences between ER subtypes (F _(2,45)_ = 180.3; p < 0.0001, n^2^ = 0.889). A Bonferroni multiple comparisons test determined a significant difference in relative expression of GPER1 versus ERα (p < 0.0001) and GPER1 versus ERβ (p < 0.0001) but no difference between ERα and ERβ (p > 0.9999).

## Discussion

Our study is the first to report that GPER1 activation in the DLS modulates preference formation for cocaine in male rats. We have shown that activation of GPER1 in the DLS is sufficient to attenuate cocaine CPP, while inhibition of GPER1 receptors produces a significant preference for cocaine at a dose that does not usually induce CPP in males. We also identified the relative mRNA expression of ERs in the dorsal striatum and found no sex differences, indicating that the sex-specific effect of GPER1 activation/inhibition is not due to an overall sex difference in the expression of receptors within this brain region.

Previous work in male rats found that animals that formed a strong taste aversion to amphetamine, relative to animals that formed a low taste aversion, also formed a stronger place preference for amphetamine, suggesting a common mechanism mediates both effects ^29^. Our findings that GPER1 activation inhibits preference for cocaine in males was replicable with an alternative reward, 0.1% SACC, which was preferred in control males as well as both experimental and control females. These animals still drank comparable amounts of liquid and consumed equitable amounts of chow compared to controls, which suggests that GPER1 activation does not cause an overall malaise in male animals. Instead, we hypothesize that GPER1 activation could be altering the learned rewarding effects of cocaine and SACC. In both experiments, the rewarding stimuli were introduced after G1 was administered into the DLS. Future experiments are needed to determine whether GPER1 activation would decrease the rewarding properties of cocaine or SACC after a preference has been established, or if the effect of GPER1 is limited to the initial establishment of a preference.

We initially hypothesized that the pharmacological manipulation of ERs in the DLS of females would alter their cocaine CPP, based on previous findings that E2 regulates the rewarding properties of cocaine in females ^30^. We predicted that administering ICI, an ERα and ERβ antagonist would inhibit CPP formation, but our results do not support this. Both ICI and G1 attenuated cocaine CPP in males, but not females. Since ICI is an ERα and ERβ antagonist, and a GPER1 agonist, this suggests that it was the GPER1 agonist action that attenuated CPP in males. We postulate that the effects of ICI in females were seen because it is also an agonist for GPER1 and that we would have needed to give an antagonist all three receptors to inhibit cocaine preference formation in females. We also recognize that a significant limitation of this study is the lack of varied doses, as we only administered 10% drug:vehicle in both sexes. Although this was a sufficient dose in males, it could be that females need a higher or lower dose to cause a change in behavior.

Females show enhanced rotational behavior and sensitization after cocaine exposure compared to males and this effect is mediated by ERα ^25,31^. Work in dissociated medium spiny neurons from dorsal striatum of females has shown E2 decreases L-type calcium current, which implicates ERβ ^32,33^. In neurons from males, the response to E2 was significantly less ^34^. Finally, E2 treatment reduces GABA release, and overexpression of ERα in the dorsal striatum also enhances the inhibitory effects of E2 on GABA release ^31,35^. Thus, ERα and ERβ are implicated in the effects of E2 on striatal function in females.

Our study used qPCR to explore relative RNA expression of ERα, ERβ, and GPER1 in the dorsal striatum and we did not find any sex differences. These findings are consistent with recent evidence supporting no sex differences in protein levels of ERα or GPER1 in the dorsal striatum of adult males and females ^36^. Our study did determine that GPER1 mRNA levels are greater than ERα and ERβ, which do not differ from one another.

We did not differentiate the medial versus lateral subsections of the dorsal striatum, and distribution of ERs within the dorsal striatum could differ by sex. It is known that in females, membrane receptors that are coupled to mGluR mediate the rapid effects of E2 in the DLS, but this mechanism has not been investigated in males ^22,24^. Whether there are sex differences in the signaling pathways mediating the effects of E2 in the DLS needs to be explored. Together, these data suggest that the sex-specific behavioral outcomes of GPER1 manipulation are likely due to differences in the downstream effects of receptor activation, rather than sex differences in overall expression.

In conclusion, we report that we have identified a novel role for GPER1 in males. To our knowledge, this is the first set of studies to show that activation or inhibition of GPER1 in the DLS is sufficient to alter cocaine conditioned place preference in males. Based on the role of E2 seen in female drug-seeking, we historically hypothesized an increase in motivated behaviors behaviors after E2 treatment, and therefore designed studies to detect an increase in drug-seeking or drug preference, rather than a decrease. This could be one reason that a role for E2 in drug-seeking in males has been missed until now. Given these results, we postulate that GPER1 is a potential target for decreasing motivation to attain cocaine in males, which is currently under investigation in our laboratory.

## Acknowledgements

In alphabetical order, we thank Kendra Beaudoin, Hannah Epstein, Brianna Graham, Lahin Lalani and Benjamin Lipkin for assisting in craniotomies, day-to-day animal handling as well as behavioral testing. We would also like to acknowledge Caitlin Posillico for her integral role in tissue processing and qPCR data collection. We thank Molly Logsdon for helping with creating the figures for this manuscript and Brandon Luma for the construction of the chambers used in the CPP behavioral Paradigm.

## Author Contributions

All authors had full access to all the data in the study and take responsibility for the integrity of the data and accuracy of the data analysis. *Conceptualization:* Quigley, J.A. and Becker, J.B.; *Methodology:* Quigley, J.A. and Becker, J.B.; *Investigation:* Quigley, J.A.; *Formal Analysis:* Quigley, J.A.; *Resources:* Becker, J.B.; *Writing – Original Draft:* Quigley, J.A.; *Writing – Revision and Editing:* Quigley, J.A. and Becker, J.B.; *Visualization:* Quigley, J.A.; *Supervision:* Becker, J.B.; *Funding:* Becker, J.B.

